# NeuronBridge: an intuitive web application for neuronal morphology search across large data sets

**DOI:** 10.1101/2022.07.20.500311

**Authors:** Jody Clements, Cristian Goina, Philip M. Hubbard, Takashi Kawase, Donald J. Olbris, Hideo Otsuna, Robert Svirskas, Konrad Rokicki

## Abstract

Neuroscience research in *Drosophila* is benefiting from large-scale connectomics efforts using electron microscopy (EM) to reveal all the neurons in a brain and their connections. In order to exploit this knowledge base, researchers target individual neurons and study their function. Therefore, vast libraries of fly driver lines expressing fluorescent reporter genes in sets of neurons have been created and imaged using confocal light microscopy (LM). However, creating a fly line for driving gene expression within a single neuron found in the EM connectome remains a challenge, as it typically requires identifying a pair of fly lines where only the neuron of interest is expressed in both. This task and other emerging scientific workflows require finding similar neurons across large data sets imaged using different modalities. Here, we present NeuronBridge, a web application for easily and rapidly finding putative morphological matches between large datasets of neurons imaged using different modalities. We describe the functionality and construction of the NeuronBridge service, including its user-friendly GUI, data model, serverless cloud architecture, and massively parallel image search engine. NeuronBridge is openly accessible at http://neuronbridge.janelia.org/.

## Introduction

Driven by advances in focused ion beam scanning electron microscope (FIB-SEM) technology (Xu et al. 2017), large-scale efforts such as Janelia’s FlyEM Project are producing detailed maps, called connectomes, of the *Drosophila melanogaster* central nervous system (Scheffer et al. 2020). Connectomes derived from EM images describe the precise structure of the neurons in a brain, as well as the connections (i.e., synapses) between them. Neuroscientists use information from the connectome to drive experiments investigating how neurons function *in vivo.*

Importantly, neuron morphology is highly conserved across different specimens in *Drosophila* (Schlegel et al. 2021), enabling the study of selected neuronal circuits across individuals. Researchers target one or more carefully selected neurons for visualization, neuronal activity measurement, genetic modification, ablation, or stimulation, and combine these tools with animal behavior studies. Large driver line libraries such as Janelia’s FlyLight Generation 1 (Gen1) GAL4 Collection (Jenett et al. 2012) provide the starting point for targeting specific neurons. Each driver line expresses the GAL4 protein under control of a genomic enhancer fragment. When crossed to another line with a UAS (Upstream Activating Sequence) reporter gene, GAL4 binds to UAS and activates reporter expression in defined neurons (Brand and Perrimon 1993).

To provide greater specificity and target individual neurons, the two-component Split-GAL4 method (Luan et al. 2020; Pfeiffer et al. 2010) is used to produce a pattern reflecting only the common elements in the expression patterns of two lines; i.e., an intersection between two expression patterns is performed. Ideally, this results in a new driver line that targets a single neuron or cell type (Fig. 1). Subsequently, the GAL4/UAS system (Brand and Perrimon 1993) allows further genetic modifications of the targeted neurons (fluorescence, ablation, etc.).

**Figure 1.**
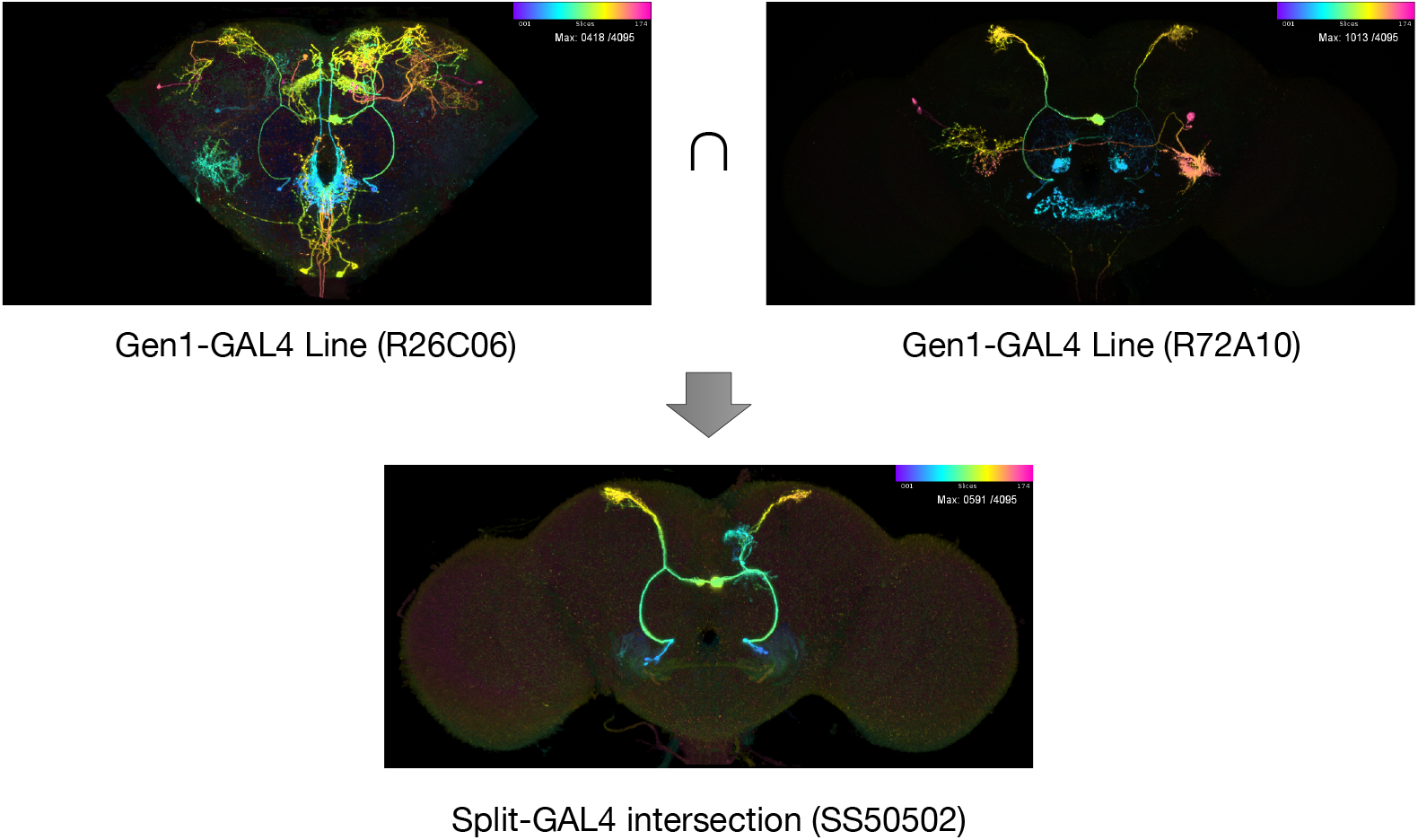
Genetic tools for *Drosophila*. Two Gen1 driver lines with broad expression are crossed using Split-GAL4 to create a new line with the intersection of the parent lines’ expression pattern. Images are color depth maximum intensity projections (CDM), where blue indicates the anterior of the brain and red the posterior. Upper section shows one of three MCFO reporter colors as a CDM. Lower section shows the full Split-GAL4 expression pattern from line SS50502 (Wolff and Rubin 2018). Original image stacks are available at https://gen1mcfo.janelia.org and https://splitgal4.janelia.org.

Creating an effective Split-GAL4 line requires the careful identification of two parent Gen1 GAL4 lines which have expression in the same neuron of interest with no other expression in common. Finding such candidate lines in a collection of thousands of images is a labor-intensive process and benefits from a computationally assisted workflow (Otsuna, Ito, and Kawase 2018). The FlyLight Gen1 GAL4 driver collection has been characterized recently (Meissner et al. 2022) with confocal imaging of neurons isolated by stochastic multi-color labeling (MCFO; Nern, Pfeiffer, and Rubin 2015). This allows for the visualization and identification of individual neurons and provides a basis for a computational workflow for Split-GAL4 creation (Meissner et al. 2022).

Increasing numbers of neuron and cell-type specific lines are being characterized (Gao et al. 2008; Tuthill et al. 2013; Aso, Hattori, et al. 2014; Wu et al. 2016; Aso and Rubin 2016; Robie et al. 2017; Namiki, Dickinson, et al. 2017; Wolff and Rubin 2018; Dolan et al. 2019; Bidaye et al. 2020; Davis et al. 2020; Morimoto et al. 2020; Schretter et al. 2020; Feng et al. 2020; Wang et al. 2021; Sterne et al. 2021; Nojima et al. 2021; Tanaka and Clark 2022; Namiki, Ros, et al. 2022; Israel et al. 2022). As the number of published Split-GAL4 lines increases, a second workflow will become important: searching the published Split-GAL4 lines to find ready-made drivers for neurons identified in the connectome.

In other common workflows, researchers traversing circuits in the EM connectome want to verify cell type identity by referencing known fly lines (Bidaye et al. 2020; Nojima et al. 2021; Sareen, McCurdy, and Nitabach 2021), or by finding additional neurons of the same cell type with a similar morphology to a known neuron of interest (Morimoto et al. 2020). In yet another workflow, a researcher starts with a neuron imaged in LM and looks for that neuron in the EM connectome (Israel et al. 2022).

All of the above workflows require identifying similar neuron morphology across large data sets of EM and LM images. Registration of neurons to a common reference alignment space enables spatial matching across samples and modalities (Bogovic, Otsuna, Heinrich, et al. 2020). Two current solutions to the computational problem of matching neurons across registered imaging modalities are Color Depth MIP Search (abbreviated as CDM Search; Otsuna, Ito, and Kawase 2018) and PatchPerPixMatch (PPPM; Mais et al. 2021). CDM Search represents location in the projection dimension as color in a MIP (maximum intensity projection) and thus enables efficient 3D structure comparison through simple 2D similarity between overlapping pixels. PPPM leverages deep-learning segmentation (PPP; Hirsch, Mais, and Kainmueller 2020) of the LM images and an algorithm based on NBLAST (Costa et al. 2016) to find the best matching LM neuron fragments that fit an EM neuron in 3D space. A comparison of CDM Search and PPPM is available in Meissner et al. 2022.

However, the CDM and PPPM algorithms are not readily accessible to experimentalists. CDM Search was initially implemented as a Fiji plugin (Schindelin et al. 2012) for executing searches locally. Therefore, it requires the user to download large data sets and wait for each search to execute using local compute resources. PPPM provides precomputed results for a subset of Gen1 MCFO images and the hemibrain dataset, but the algorithm is expensive to run (~3 hours per LM volume on a single GPU) and is not easily usable for custom searching with user data. To make these algorithms more accessible, we built a web application that experimentalists can use to rapidly identify and view similar neurons in published EM and LM data sets, or to perform searches on their own data. The only comparable software that we know of is the NBLAST functionality in Virtual Fly Brain (Milyaev et al. 2012), which allows users to find similar neurons across the FlyCircuit data set. However, it is currently limited to EM data sets and does not address the use cases of EM-LM search.

## Results

To address the problem of finding similar neurons across large multi-modal data sets, we developed NeuronBridge, an easy-to-use web application. It provides instant access to neuron morphology matches for published EM and LM data sets, as well as rapid custom searching against those data sets. Our initial implementation was based on the CDM Search algorithm and PPPM results were added later. Implementing CDM Search in a publicly available web application with interactive response times presented several major challenges: a) the CDM Search tools were only available via a Fiji GUI and lacked programmatic APIs for reuse; b) when centralized and scaled to hundreds of users, CDM searching is expensive to run *de novo* for each query; c) CDM Search relies on fast local disk access and multithreading, which are expensive resources to leave idle at the scale needed to support unpredictable usage access patterns; and d) users seeking to search using their own data first need to align it to the standard template and generate aligned CDM images for their neurons.

We first extracted the CDM Search algorithm from its Fiji plugin, refactored it into a Java library (Software Availability), and made it available for reuse. To enable large-scale computation on a high performance computing (HPC) cluster, we used the CDM Search library to create an Apache Spark application for running the CDM Search algorithm in a distributed manner (Methods).

Second, we used our Apache Spark CDM Search implementation to precompute neuron matches between the published FlyEM and FlyLight data sets (Methods), including the FlyEM hemibrain (Scheffer et al. 2020) and the Gen1 MCFO (Meissner et al. 2022) and Split-GAL4 driver libraries (Fig. 2). The data in each image was first extracted into multiple files for improved searchability (Methods), and then each EM image was compared with all of the LM images, a total of 7.4 billion image-to-image comparisons. Matches were recorded in both directions, to allow bi-directional searching. This step obviated the need to run *de novo* searches for matches between published data sets, mitigating the ongoing costs of running the shared service, and allowing us to make the service freely available to the community.

**Figure 2.**
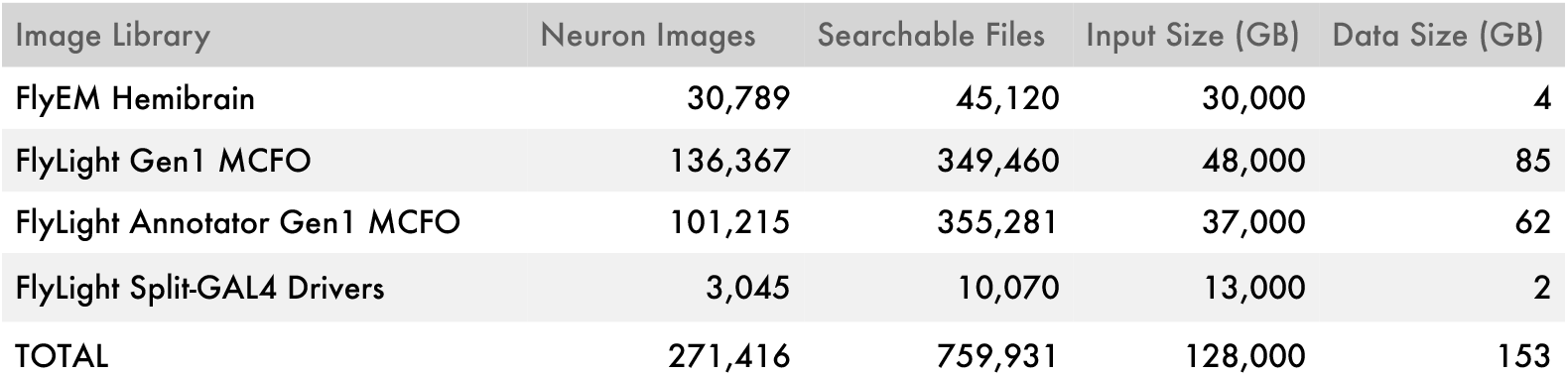
Data for which precomputed matches are available in NeuronBridge. To generate precomputed results, each image in the FlyEM hemibrain library was compared against all of the FlyLight images, and matches were recorded in both directions. *Neuron Images* indicates the total number of individual images containing neurons. *Searchable Files* indicates the number of CDM images for searching, after pre-processing steps of segmentation and flipping (Methods). *Input Size* indicates the total size of all original imagery compressed losslessly. *Data Size* indicates the total size of the CDM images used for searching.

Third, we developed a data model (Methods) for representing matches that we define as putatively similar neuronal morphologies identified with respect to two images that are registered to the same template. Our data model allows matches from CDM Search, PPPM, and potentially any future algorithms to be represented with the same entity classes (Methods), as well as allowing for an extensible set of imaging modalities, making it easier to add new data in the future while maintaining a consistent user experience.

Then we imported the published PPPM results (Mais et al. 2021) for the hemibrain (EM) and Gen1 MCFO (LM) into our data model alongside the CDM Search results. PPPM compares each EM neuron to each MCFO sample using all three channels at once, so the LM channel information for each match is left empty. In addition, the PPPM results include additional result images for visualizing matches, including overlays of EM skeletons on top of masked LM signals, which we were able to import into our data model.

Next, we leveraged the Amazon Web Services (AWS) Open Data program to make all of the images and match metadata available as a robust public data API (Methods) on AWS S3. AWS S3 is a serverless object storage service with high reliability, low latency, access control, and tight integration with other AWS services. Using a serverless solution allowed us to keep costs low compared to a traditional relational database deployment. We also developed a simple Python module (Software Availability) on top of our API, to enable ad hoc data analysis through the use of Python libraries, and to serve as a client reference implementation.

Next, we created an online custom search service for user-uploaded data (Methods) that 1) registers uploaded confocal image stacks to a reference brain using CMTK (Rohlfing and Maurer 2003) running on AWS Batch, 2) generates a CDM image for each channel of the uploaded stack, 3) allows the user to select their neurons of interest with an interactive browser-based image masking tool, and 4) runs a massively parallel CDM Search to find neuron matches in the EM. Importantly, this component is entirely serverless, keeping costs low by not idling compute resources in the cloud. To make this possible we developed a burst-parallel compute engine (Methods) to achieve on-demand, 3000-fold parallelism using the AWS Lambda service, which allows a search of the EM data set to complete in seconds and a search of the LM data sets to complete in under two minutes.

Finally, we built a user-friendly, single-page application (SPA) using the React framework (Methods), enabling end-users to browse precomputed matches and run searches with their own data. The GUI of the application (Fig. 3) was designed to be simple and intuitive, focusing solely on the task of neuron matching, providing contextual help information, and including usage examples where appropriate.

**Figure 3.**
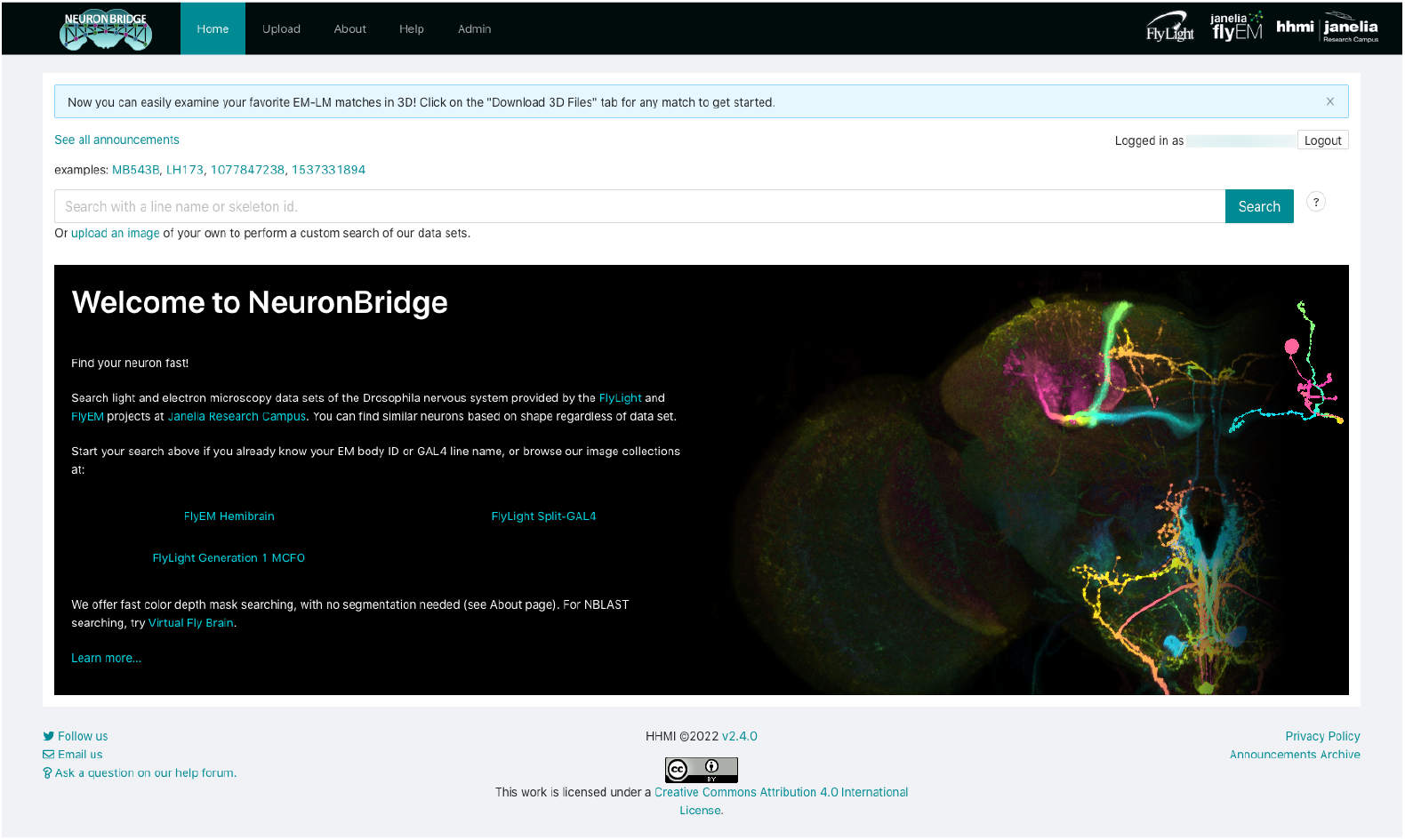
NeuronBridge home page. Google-like search box allows rapid look-up of EM and LM images via common identifiers such as body ids and published line names.

NeuronBridge currently serves over 150 million precomputed matches between 5,860 fly lines and 30,789 EM neurons. In addition, these lines and neurons have been made available for online custom searching, comprising a data set of 759,931 CDM images. These searchable 2D images efficiently represent signals extracted from approximately 98 TB of confocal light microscopy and 30 TB of electron microscopy data (Fig. 2) in 150 GB of searchable images, a three orders of magnitude compression which operationalizes the data for neuron searching. We additionally provide aligned EM neurons in SWC format and aligned 3D stacks for LM images in H5J format (https://data.janelia.org/h5j), so that putative matches can be validated manually in 3D using an external system like VVD Viewer (Kawase and Otsuna 2022), to supplement the 3D viewing supported directly in the GUI (3D Visualization).

## Discussion

All data served by NeuronBridge is available for programmatic access via open data APIs (Data Availability) allowing for reuse and integration. These APIs have already enabled several third-party applications including the Neuronbridger R API (http://natverse.org/neuronbridger) and FlyBrainLab NeuroNLP (Lazar et al. 2021). The serverless back-end automatically scales to meet additional load from these API users and imposes rate limits on users who adversely affect service performance.

In the future, NeuronBridge could benefit from improved and/or new neuron matching algorithms, additional functionality for match inspection, and any other improvements that address match quality or the core workflows of match browsing, verification, and export. We expect this resource to grow with additional data, including additional EM volumes expanding the connectome coverage of the fly nervous system, as well as images of new Split-GAL4 lines.

We intend for NeuronBridge to remain a very focused tool, and we have therefore intentionally constrained its architecture and implementation (Methods) to solving the neuron matching use cases. We provide contextual links in cases where other web applications already provide related functionality. For access to source LM images, we link to FlyLight’s anatomy websites (https://flweb.janelia.org, https://gen1mcfo.janelia.org, https://splitgal4.janelia.org). For EM neurons we link to neuPrint (Clements et al. 2020, https://neuprint.janelia.org). We also provide cross reference links to the Virtual Fly Brain (Milyaev et al. 2012) for both EM and LM results. All of these websites link back to NeuronBridge, forming a synergistic ecosystem of tools and data.

We hope that the cloud-based serverless architecture of NeuronBridge, uncommon in open source scientific research software, inspires architectural decisions or code reuse in other tools and platforms for large scientific data analysis. In particular, the burst-parallel image search balances large data analysis with online query capability on interactive timescales, and could be reused to perform many other types of analysis where a large data volume must be fully traversed while a user is waiting.

The NeuronBridge web application fills a gap created by recently published large data sets in *Drosophila* neuroscience including the EM connectome and LM images for characterizing driver lines. Researchers making use of the connectome can now efficiently find fly lines that target their neurons of interest, as well as search these data sets to verify cell type identity and driver line expression. We believe that these properties will make NeuronBridge an indispensable tool for *Drosophila* neuroscience research.

## Methods

### Image Alignment

For the EM hemibrain data set we relied on the published registration to JRC2018 (Bogovic, Otsuna, Heinrich, et al. 2020) (Fig. 4, D). We used the Computational Morphometry Toolkit (CMTK, Rohlfing and Maurer 2003) to register all LM brains (Fig. 5, A-B) to the unbiased gender-specific JRC2018 templates (Bogovic, Otsuna, Heinrich, et al. 2020). In both cases we subsequently used a bridging transform to move the images into the alignment space of the Unisex template, where neuron matching could be performed irrespective of gender.

**Figure 4.**
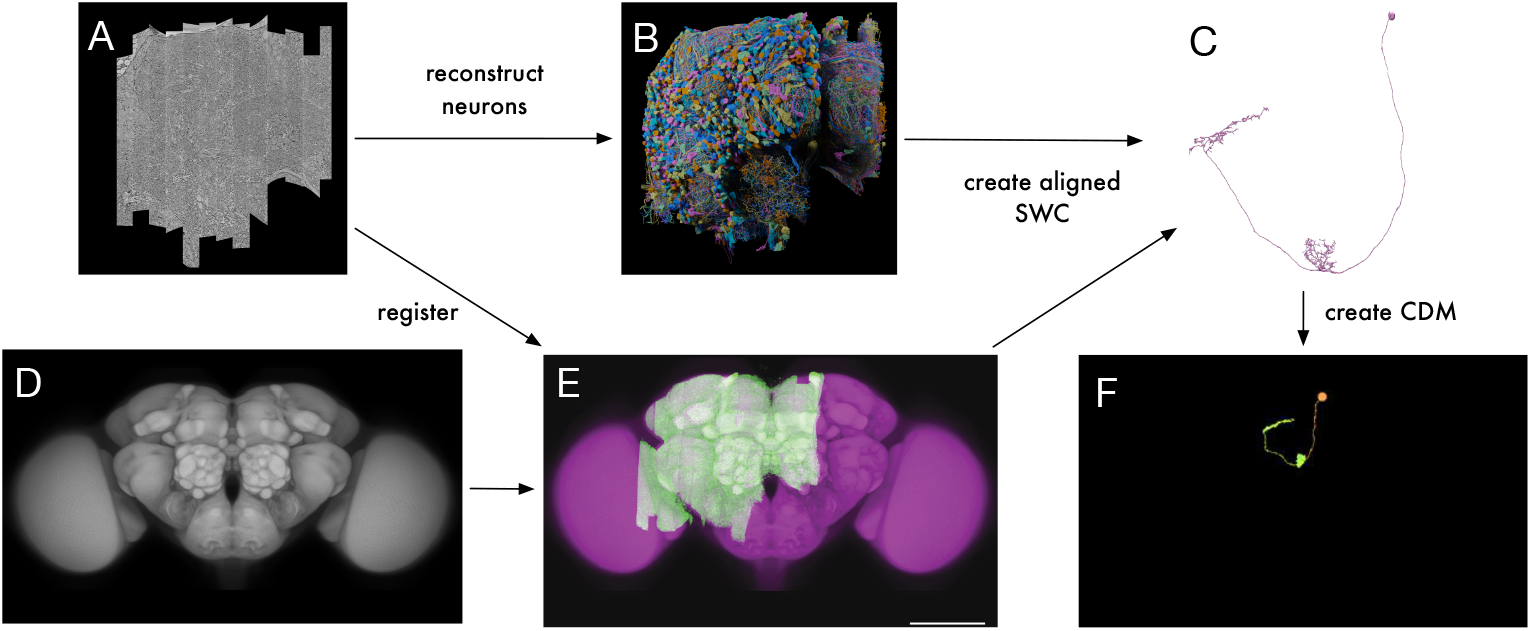
EM pipeline for CDM generation. (A) FlyEM hemibrain stitched data; screenshot from Neuroglancer (B) Hemibrain reconstruction by Janelia FlyEM and Google (C) Example hemibrain neuron with body ID 1537331894; screenshot from neuPrint (D) JRC2018_Unisex_HR brain template (E) Registration of hemibrain to the JRC2018_Unisex template (F) CDM of the neuron shown in C.

**Figure 5.**
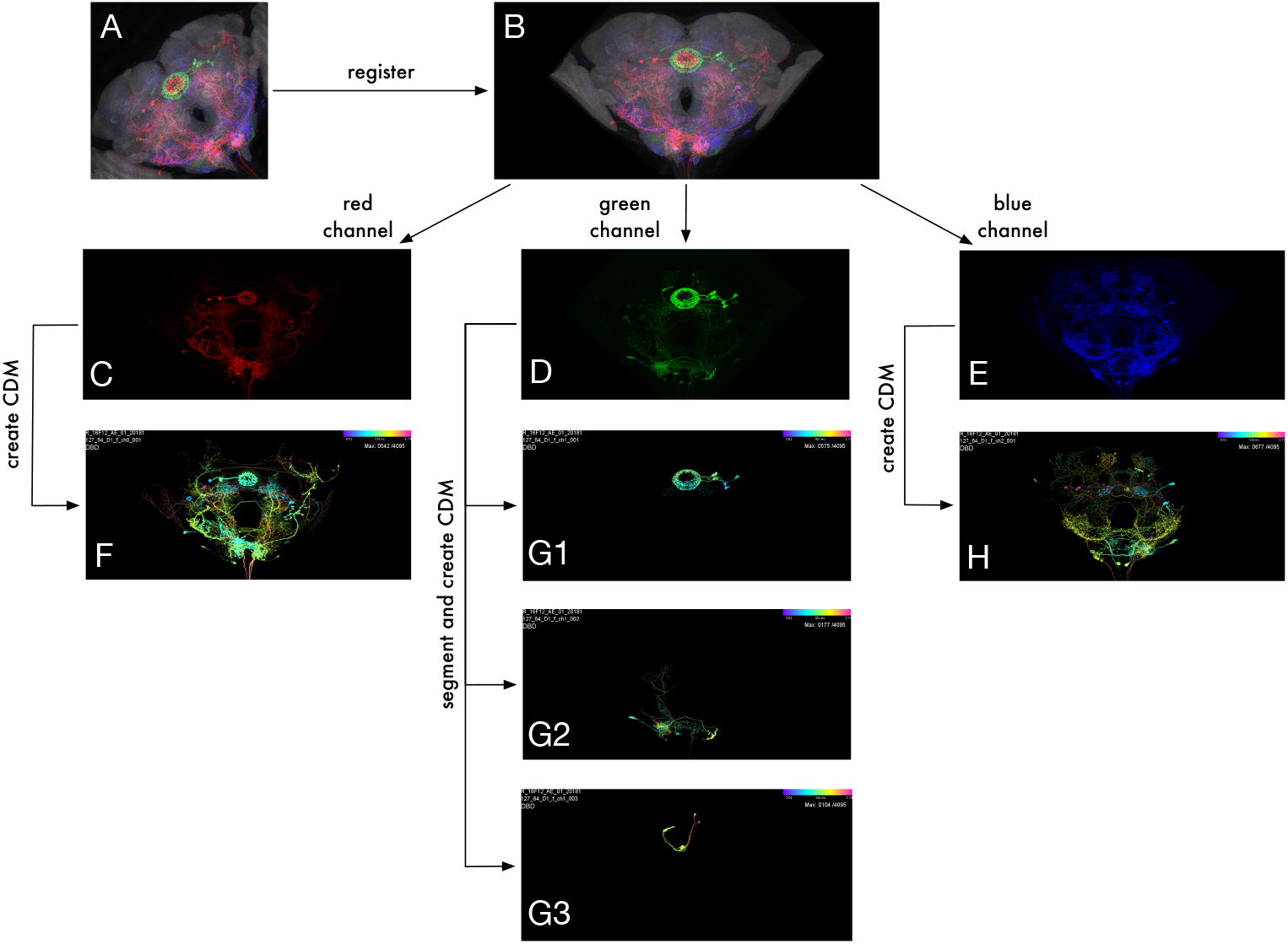
LM data pipeline. (A) Example LM MCFO image of R16F12 line with slide code 20181127_64_D1 (B) LM image registration (C-E) Aligned MIPs of individual channels (F-H) CDM images for each channel (G1-G3) Segmentation of green channel using modified DSLT yields three voxel sets for the green channel.

For online searching, the alignment code is packaged as a Docker container and run using AWS Batch (Software Availability). For better performance, online searches skip the gender-specific alignment and align directly to the Unisex template.

### Neuron Morphology Matching

The CDM Search algorithm (Otsuna, Ito, and Kawase 2018) was designed for interactive use and required several modifications for reliable batch execution on large data sets (Meissner et al. 2022). These changes will be described fully in a future paper (Bogovic, Otsuna, Takemura, et al. 2022) but we provide a brief outline here. First, to reduce occlusions, especially in denser MCFO samples, we performed a 3D segmentation of all LM images using an algorithm based on DSLT (Kawase, Sugano, et al. 2015), which resulted in a handful of voxel sets per image channel (Fig. 5, G). We eliminated smaller “junk” voxel sets based on manually chosen Fiji shape descriptor thresholds and generated CDM images of the remaining voxel subsets.

As its name suggests, the EM hemibrain does not cover the entire fly brain, since it is assumed that many of the neurons exhibit bilateral symmetry (Scheffer et al. 2020). To improve precomputed matches, we created additional artificial CDM images by mirroring any EM neurons that crossed the midline. These “flipped” neurons augment the EM image library by making it more likely for the algorithm to find bilateral neurons which appear on both sides of the brain in LM images.

Next, we modified the CDM Search algorithm to penalize expression in one image that is not present in the other, to devalue matches which share matching pixels but also have large areas of non-coincidental expression. The final score for each match was computed as a ratio of positive matching (i.e., number of matched pixels) to negative matching (i.e., non-coincidental expression). When running EM/LM search, we used EM single neuron bodies as search masks and compared them with every LM image. Every result was recorded as a match in both directions, EM→LM and LM→EM. For EM→LM, we limited the number of matches to a maximum of 300 lines and 3 matches per line.

We also encapsulated the CDM Search algorithm into a reusable Java library and used that library in an Apache Spark application which scales computation on an HPC cluster. Finally, we translated the core CDM Search algorithm into JavaScript, to reduce cold starts when running searches on AWS Lambda.

### User Experience

NeuronBridge is an end-user web application designed for accessibility and ease-of-use by wet-lab experimentalists. Using any web browser, a user can log into the service’s website with their Google account credentials or create a separate NeuronBridge account. The user is presented with a simple search interface for looking up their neuron body or line of interest (Fig. 3). Searching for an EM body returns a CDM representation of the body, in the common alignment space. Searching by neuron name may return multiple EM bodies to choose from. Searching for an LM line may likewise return multiple images, since the line may have been imaged multiple times, with each having multiple color channels. These initial search results are displayed in a tabular, paginated format, allowing the user to select an image of interest to begin neuron matching. Each result has buttons corresponding to the types of match algorithms that were used to compare it to the other modality, typically both CDM Search and PatchPerPixMatch. These buttons let the user view putative neuron matches with other images (LM matches for EM targets, EM matches for LM targets).

After selecting a target image and an algorithm, precomputed neuron matches are immediately displayed as a list or a grid (Fig. 6) and paginated for rapid browsing. They can be filtered by various criteria, such as library, body ID, or line name. When searching for LM matches, all of the matches for a line are grouped and sorted together by the score of the highest ranking representative of that line. These line results can be filtered to display a maximum number of representatives per line. A setting of 1 is useful when the user is interested in fly lines as the final output.

**Figure 6.**
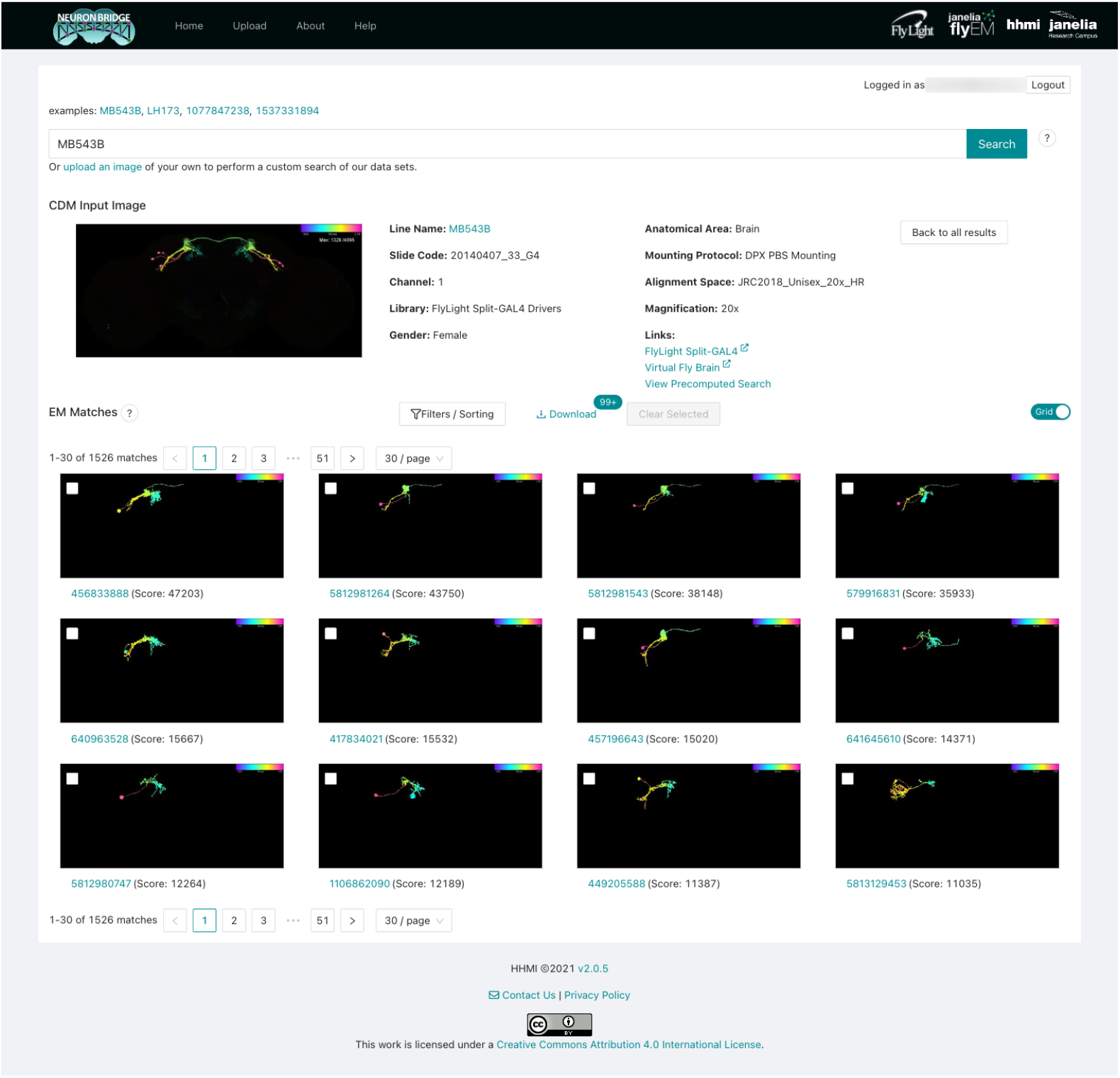
Browsing EM matches for an LM image. Paginated EM matches are displayed for the CDM input image shown at the top. The results can be filtered and sorted, or shown as a list. Checkboxes allow users to select promising matches for download and further analysis. Screenshot was truncated to only show the first three rows of results.

Clicking one of the matches brings up a detail interface (Fig. 12, Appendix) which allows the user to compare the match to the target using a synchronized cursor. Various accessory images can be displayed to better characterize the match and the fly line or EM volume where it was found. Matches can be selected for download, from either the overview page or the detail page. The user can download all the metadata for the selected matches as a CSV file, and all of the images as a ZIP file.

In addition to the precomputed match results between published images, NeuronBridge supports custom CDM Search against any of the EM and LM image libraries. Users begin a custom search by uploading their own image (Fig. 16, Appendix), which can be an unaligned image stack in a variety of standard formats (TIFF, ZIP, LSM, OIB, CZI, ND2), or an aligned CDM mask in the JRC2018_Unisex_HR alignment space. NeuronBridge attempts to align uploaded image stacks and generates a CDM for each channel. This process may take several minutes, and the user may wait for the alignment task to complete, or come back later to find their aligned CDMs. Next, the user is asked to choose a channel and create a search mask. The user may select either EM or LM libraries to search, and optionally specify CDM Search parameters before invoking the search. A custom search typically takes less than a minute to process and progress is displayed in a step-wise workflow GUI (Fig. 7). Results are presented identically to how precomputed results are displayed.

**Figure 7.**
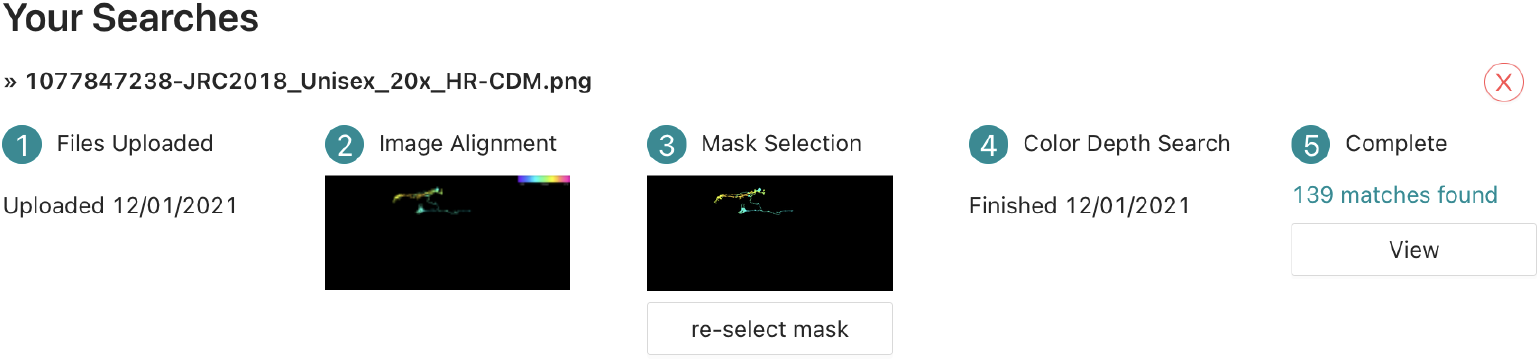
Custom search progress steps. The workflow can be branched by re-selecting the mask so that a different search can be run with the same input stack. This saves a considerable amount of time by reusing the alignment result.

The web GUI was developed using the React framework in order to make the GUI components reusable and more responsive than a traditional web application, by never fully reloading the page while moving around the site. Instead, data is loaded via asynchronous HTTP requests, and the GUI is updated as data becomes available. By using AWS AppSync, we also extended this idea to asynchronous updates using Web Sockets, so that long-running back-end operations such as alignments can be monitored without polling. The single-page application (SPA) approach also allowed us to serve the website as static content, with caching via the AWS CloudFront Content Distribution Network (CDN) that speeds up initial loading.

### 3D Visualization

NeuronBridge provides two ways a user can manually verify a putative match by viewing the EM neurons and LM images from arbitrary viewpoints in 3D. The simplest approach is to click the “View in 3D” button in the detail interface for a search result (Fig. 12, Appendix). Doing so launches a new browser page that loads the LM image volume and EM neuron skeleton and renders them in 3D with interactive camera controls (Fig. 14, Appendix). For the LM images we use direct volume rendering, which samples the image volume along rays from the camera, applies a transfer function and lighting model to compute color and opacity at each sample, and composites the results to produce pixels. The EM neuron is rendered as an opaque surface and blended into the volume rendering with proper occlusion based on its depth map. The renderer uses WebGL2 and achieves interactive performance on modern desktop and laptop computers. The URL for the renderer page is continually updated to encode the details of the current view, so a live rendering can be shared with a collaborator by sharing the URL.

A more sophisticated rendering of the EM and LM data is available by clicking the “Download 3D Files” button and loading the data into a desktop rendering application like VVD Viewer (Kawase and Otsuna 2022) (Fig. 15, Appendix). VVD Viewer renders with slightly higher quality because it uses more bits per voxel of the LM data (12 instead of 8). It also adds several capabilities beyond the browser-based renderer. It can render multiple channels of an LM dataset simultaneously, better representing the data used by the PPPM matching algorithm. It can render multiple EM bodies simultaneously, which can be useful for verifying fragmented bodies or for comparing several match results at once. It can also generate animated videos, showing changes to the camera position and other viewing parameters.

### Software Architecture

We used fully-managed, serverless AWS services to build the back-end (Fig. 8), including Cognito identity federation for authentication, IAM for authorization, Lambda for compute, S3 for storage, and DynamoDB as the database. We used only serverless services to keep operational costs low while allowing us to focus on application logic instead of spending time on server administration.

**Figure 8.**
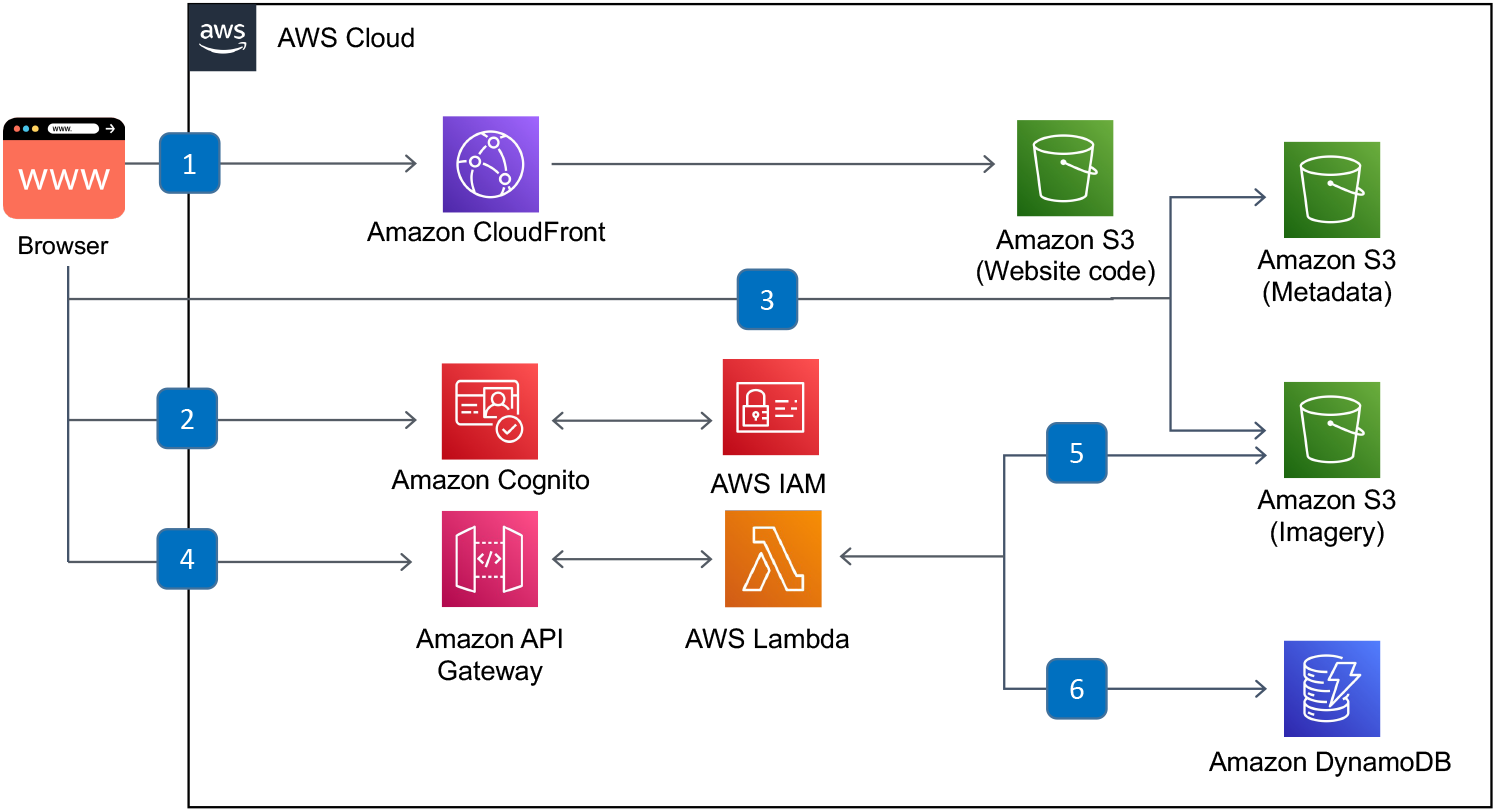
Serverless back-end for precomputed result browsing. The user’s browser communicates with (1) the CloudFront Content Distribution Network (CDN) to retrieve the web client code which is cached from AWS S3. (2) Authentication is done using federated identity providers through Cognito, supporting both email-based accounts and Google accounts, and delegating to IAM for authorization. Precomputed results are (3) statically loaded from public buckets on S3. Dynamic features use (4) an API Gateway endpoint to run a Lambda function, for example to (5) create a ZIP archive of multiple files for download or (6) run prefix searches on DynamoDB.

Custom search (Fig. 9) consists of two services: image alignment which runs on AWS Batch, and distributed CDM Search which runs on AWS Lambda. The aligner runs on AWS Batch as a Docker container and is monitored by a Step Function which notifies the client when the alignment is completed. To enable efficient searching across large image data sets while keeping costs low, we implemented a burst-parallel (Fouladi et al. 2019) search engine built on Lambda and Step Functions (Rokicki 2021). The serverless model is a particularly good fit for the burst-parallel implementation for several additional reasons. First, massive parallelism is required to quickly search large image libraries, and the ideal level of parallelism is prohibitively expensive in a traditional server-based architecture. The 3000-fold parallelism we achieved with AWS Lambda for a few cents would require one hundred 30-core dedicated servers in a traditional data center deployment. Second, the service has unpredictable usage and can sit idle for extended periods, while at other times has bursts of high activity. This type of usage pattern is directly addressed by a serverless model that can scale to zero. The costs to run this service are low because serverless services bill for compute/storage that is utilized, similar in concept to a traditional HPC compute cluster. In addition, the serverless approach reduced the amount of ongoing maintenance by eliminating server administration. Finally, using serverless Lambda functions allowed us to fully leverage the concurrent throughput of the S3 buckets hosting the images.

**Figure 9.**
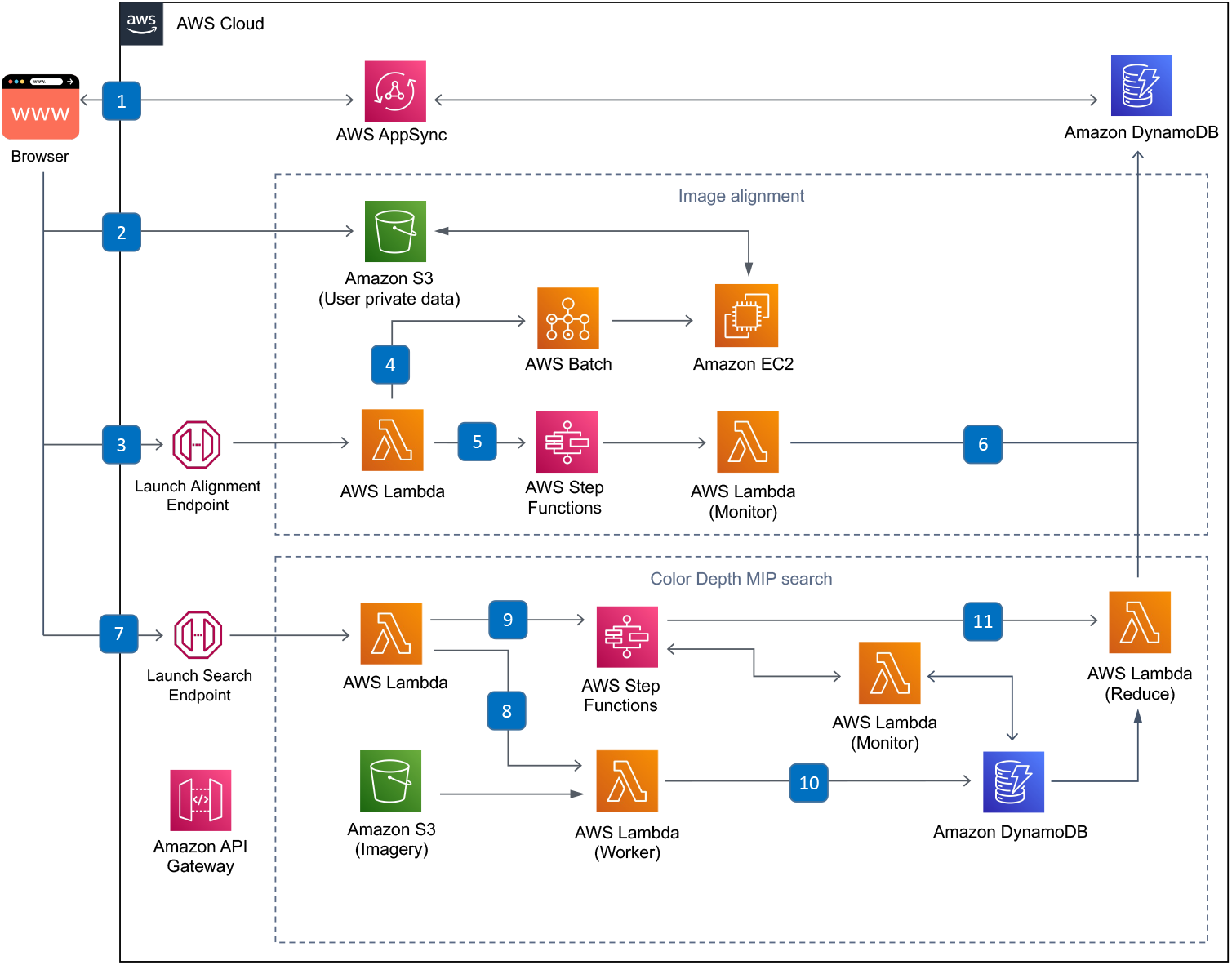
Serverless back-end for custom search. The user initiates a search by uploading an image, and three messages are sent to the back-end: (1) Using AppSync, a new record is created in DynamoDB to track the search workflow, (2) the image is uploaded to a private user folder on S3, and (3) the client starts the alignment process by invoking a Lambda function. (4) A Batch job is created, which allocates an EC2 node and runs the alignment Docker container, fetching the uploaded file from S3, and depositing the aligned CDM images back on S3. (5) A Step Function periodically runs a Lambda to monitor the progress of the Batch job. (6) Once the alignment job is finished, it updates the workflow state in DynamoDB, which notifies the user via AppSync. Back in the browser, the user is asked to create a search mask from one of the resulting CDM images, and this mask is uploaded to S3. (7) The client calls a second endpoint to begin the search, which calls a Lambda that (8) recursively starts the burst-parallel search workers as well as (9) the monitoring Step Function state machine. (10) When a worker finishes, it records its result in a DynamoDB table. Once the monitoring Lambda detects that all workers are finished, (11) it runs the reducer which produces a search result that is updated in the DynamoDB workflow table, notifying the user again through AppSync that the search is complete.

The software is reliably deployed through the use of the Serverless Framework, which generates low-level AWS CloudFormation instructions for deployment. Reproducibility and isolation of the deployment is important to being able to run multiple versions of the system at the same time. In a single AWS account, we run a separate instance for each developer, a validation instance for testing before production deployments, a pre-publication instance for internal use, and the production instance that is publicly accessible.

### Data Model

The data model (Fig. 10) is anchored by a NeuronImage, representing a set of neurons in an alignment space that can be compared with other images in that alignment space, and a set of Matches between NeuronImages, generated by either CDS or PPPM. Image metadata is denormalized wherever an image is referenced, to allow clients to fetch a single JSON file instead of making multiple requests. The DataConfig provides summary information, such as the list of possible anatomical regions, as well as providing constants which allow long, common values to be interpolated into JSON values as shorter keys, reducing disk space usage and transfer time. Importantly, this model is trivially extensible with new matching algorithms and new imaging modalities.

**Figure 10.**
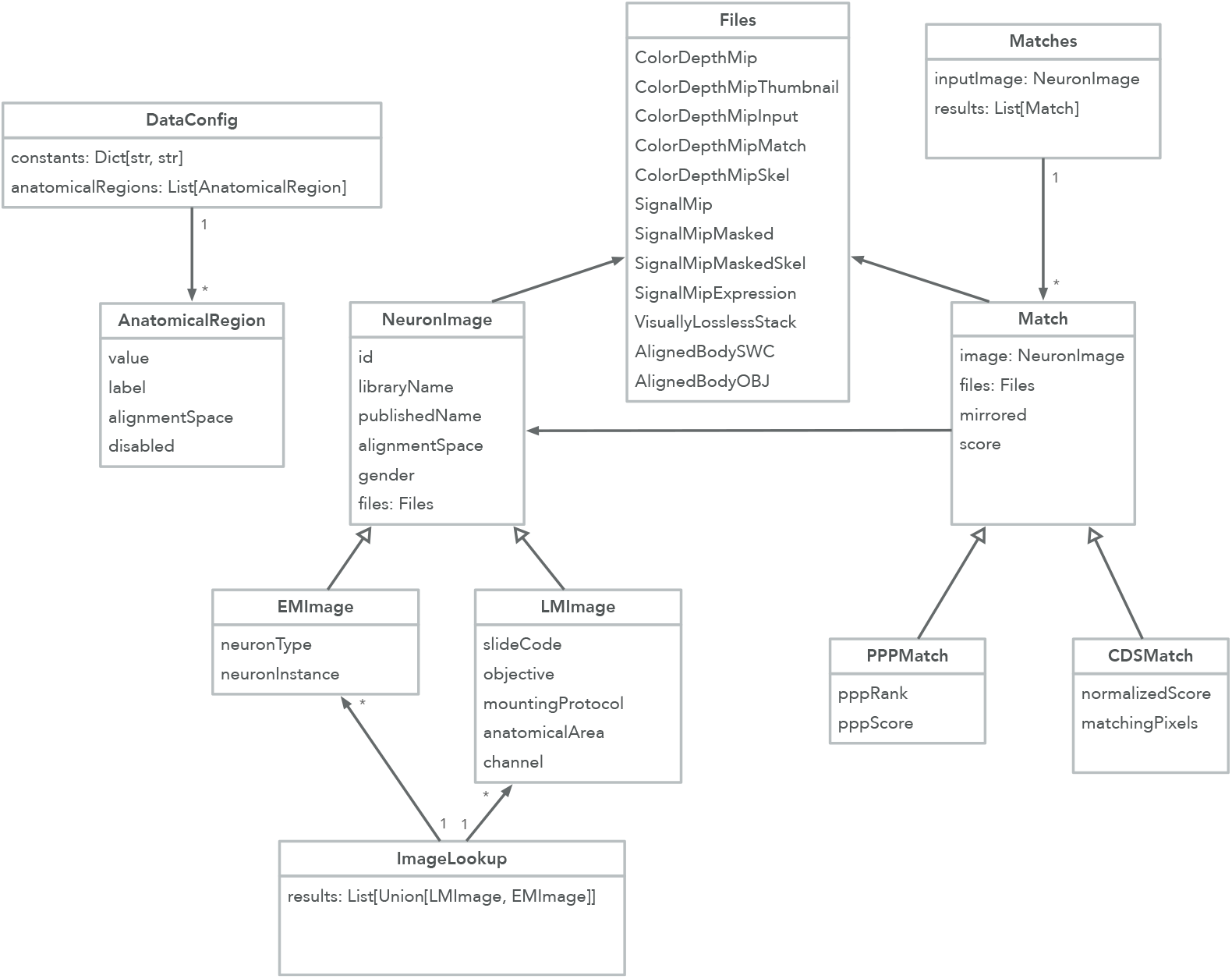
Data model. The DataConfig describes the metadata accessible through the API. Image lookup queries return an ImageLookup object containing neuron images. A NeuronImage has a GUID called id which can be queried for Matches.

### Known Limitations

We intentionally focused NeuronBridge on the use case of finding similar neuron morphologies, so it contains no features allowing users to search or view other types of data such as neuronal circuits, neurotransmitter information, or detailed metadata. The application currently only allows searching of neurons within the *Drosophila* brain, but support for the ventral nerve cord (VNC) is planned.

The quality of the matches presented by NeuronBridge is bounded by the effectiveness underlying matching algorithms. The CDM Search and PPPM algorithms both produce useful and often complementary sets of putative matches (Meissner et al. 2022). CDM Search struggles with occlusions in the projection dimension, while PPPM has difficulty with segmentation of dense samples, which can cause false positives during search (Mais et al. 2021). PPPM results are currently only available for a subset of Gen1 MCFO samples and exclude Split-GAL4, as well as the more recently-added 20x/63x Annotator MCFO data set. NeuronBridge also does not currently load reverse matches (LM→EM) for PPPM.

The architecture is scalable but limits some of the functionality in ways we anticipated. The choice of using S3 as a match database limits the queries that can be done to lookups by identifier, so users must rely on other websites for metadata-based search. Also, implementing the custom search on top of AWS Lambda limits us to a 15 minute execution time per function, though that can be worked around by using a smaller batch size and launching more Lambda functions.

### Data Availability

All public data in NeuronBridge (i.e., excluding data uploaded by users) is available for access on AWS S3. In addition to acting as a database back-end for the NeuronBridge browser application, the data on S3 serves as an Open Data API (Appendix). The data is available in the following public S3 buckets:

- Metadata `s3://janelia-neuronbridge-data-prod`

- Images `s3://janelia-flylight-imagery` `s3://janelia-flylight-color-depth` `s3://janelia-ppp-match-prod`

All of the data is licensed under the Creative Commons Attribution 4.0 International (CC BY 4.0) license.

### Software Availability

NeuronBridge is open source software and the complete source code and documentation for all of the components is available on GitHub. All of the code is licensed under a permissive Open Source license.

- NeuronBridge production instance
  – Web URL: http://neuronbridge.janelia.org
- Web client implementation
  – Source code repository: https://github.com/JaneliaSciComp/neuronbridge
  – Archived code at time of publication: 10.5281/zenodo.6702817
- back-end implementation
  – Source code repository: https://github.com/JaneliaSciComp/neuronbridge-services
  – Archived code at time of publication: 10.5281/zenodo.6588117
- Color Depth MIP search library and Apache Spark implementation
  – Source code repository: https://github.com/JaneliaSciComp/colormipsearch
  – Archived code at time of publication: 10.5281/zenodo.6588149
- Scripts for precomputation of matches
  – Source code repository: https://github.com/JaneliaSciComp/neuronbridge-precompute
  – Archived code at time of publication: 10.5281/zenodo.6588156
- Aligner implementation
  – Source code repository: https://github.com/JaneliaSciComp/neuronbridge-aligners
  – Archived code at time of publication: 10.5281/zenodo.6588136
- Utilities for uploading files to AWS and generating the DynamoDB table
  – Source code repository: https://github.com/JaneliaSciComp/neuronbridge-utilities
  – Archived code at time of publication: 10.5281/zenodo.6588121
- Python API
  – Source code repository: https://github.com/JaneliaSciComp/neuronbridge-python
  – Archived code at time of publication: 10.5281/zenodo.6486428
- 3D visualization
  – Source code repositories: https://github.com/JaneliaSciComp/neuronbridge-vol-viewer https://github.com/JaneliaSciComp/web-vol-viewer https://github.com/JaneliaSciComp/web-h5j-loader
  – Archived code at time of publication: 10.5281/zenodo.6702084, 10.5281/zenodo.6772513, 10.5281/zenodo.6772506

## Acknowledgements

We are sincerely grateful to Geoffrey Meissner for considerable input into both the NeuronBridge application design and the draft manuscript. We are likewise very grateful to Stephan Preibisch for considerable input in reviewing and editing of the draft manuscript. We would like to thank the FlyLight and FlyEM project teams for our long-term collaborations without which NeuronBridge would not have been possible. Thank you to Wyatt Korff, Reed George, and Gerry Rubin for supporting this project. Thanks to Stephen Plaza for proposing the initial idea. Thanks to Dagmar Kainmueller and Lisa Mais for help with integrating their PPPM result data set. Thanks to Scott Glasser and Ray Chang for valuable advice and prototyping of the burst-parallel search implementation on AWS. Thanks to Antje Kazimiers for early prototyping work on the web GUI. Special thanks to Brianna Yarbrough and Yisheng He for designing the NeuronBridge logo.

This software makes use of the Computational Morphometry Toolkit (http://www.nitrc.org/projects/cmtk), supported by the National Institute of Biomedical Imaging and Bioengineering. Cloud storage for images was generously provided by Amazon Web Services (AWS) as part of the AWS Open Data Sponsorship Program. The software architecture diagrams use icons from AWS Architecture Icons and flaticon.com.

Funding was provided by Howard Hughes Medical Institute. This article is subject to HHMI’s Open Access to Publications policy. HHMI lab heads and project team leads have previously granted a nonexclusive CC BY 4.0 license to the public and a sublicensable license to HHMI in their research articles. Pursuant to those licenses, the author-accepted manuscript of this article can be made freely available under a CC BY 4.0 license immediately upon publication.

## Author Contributions

J.C. designed and implemented the front-end web client. C.G. and D.J.O. implemented the back-end data processing system and generated the precomputed matches. C.G. and J.C. implemented the serverless back-end. H.O. designed and optimized the modified CDM Search algorithm and created the image alignment tools. T.K. implemented the CDM Search algorithms and optimized the custom search. R.S. uploaded and managed data on S3, and created the DynamoDB database for lookups. P.M.H. implemented the browser-based 3D visualization. K.R. conceptualized the project, designed the software architecture, managed the engineering effort, implemented the burst-parallel search framework, and wrote the draft manuscript. All authors reviewed and approved the final version of the manuscript.

## Appendix

### Open Data API

The S3 bucket containing the NeuronBridge matches forms an open RESTful API. Key structures are arranged such that they can be queried predictably. We use the standard JSON format and publish a schema for each document type. The schemas are versioned so that future changes are not breaking. All of the endpoints below are relative to the bucket root: `s3://janelia-neuronbridge-data-prod`

- `current.txt` Returns the current version, to be used in subsequent API calls as <`version`>. Older versions of the metadata are preserved so a client can choose to request older data for compatibility or other reasons.
- `<version>/schemas/` A prefix containing the version-specific JSON schemas for all objects in the data model.
- `<version>/config.json` Returns a configuration dictionary containing base URL prefixes and other metadata necessary for programmatic use of the following API.
- `<version>/DATA_NOTES.md` Returns a Markdown document containing the release notes for changes to the data from the previous version.
- `<version>/metadata/by_body/<body_id>.json` For a given EM body ID returns metadata such as a path to a representative image, neuron names, etc.
- `<version>/metadata/by_line/<line_id>.json` For a given LM fly line, returns metadata including a list of images from the line (most lines have been characterized multiple times), representative images, and other attributes.
- `<version>/metadata/cdsresults/<image_guid>.json` For a CDM neuron image provided by either the by_body or by_line endpoints, this returns a list of matching images as computed by CDM Search. Neuron images have Globally Unique Identifiers (GUID) which persist across data versions, making it easier to link to data and results.
- `<version>/metadata/pppresults/<body_id>.json` For a given EM body ID returns a list of matching LM images as computed by PatchPerPixMatch search.

### User Survey Results

We conducted a survey to assess users’ satisfaction with the service and determine which features and improvements should be prioritized. We received 22 responses to the survey during the 2021 calendar year (Fig. 11). By far, additional data was the most important aspect for users, which led us to prioritize the addition of matches for the upcoming EM ventral nerve cord (VNC) and any LM images that are released by Janelia.

**Figure 11.**
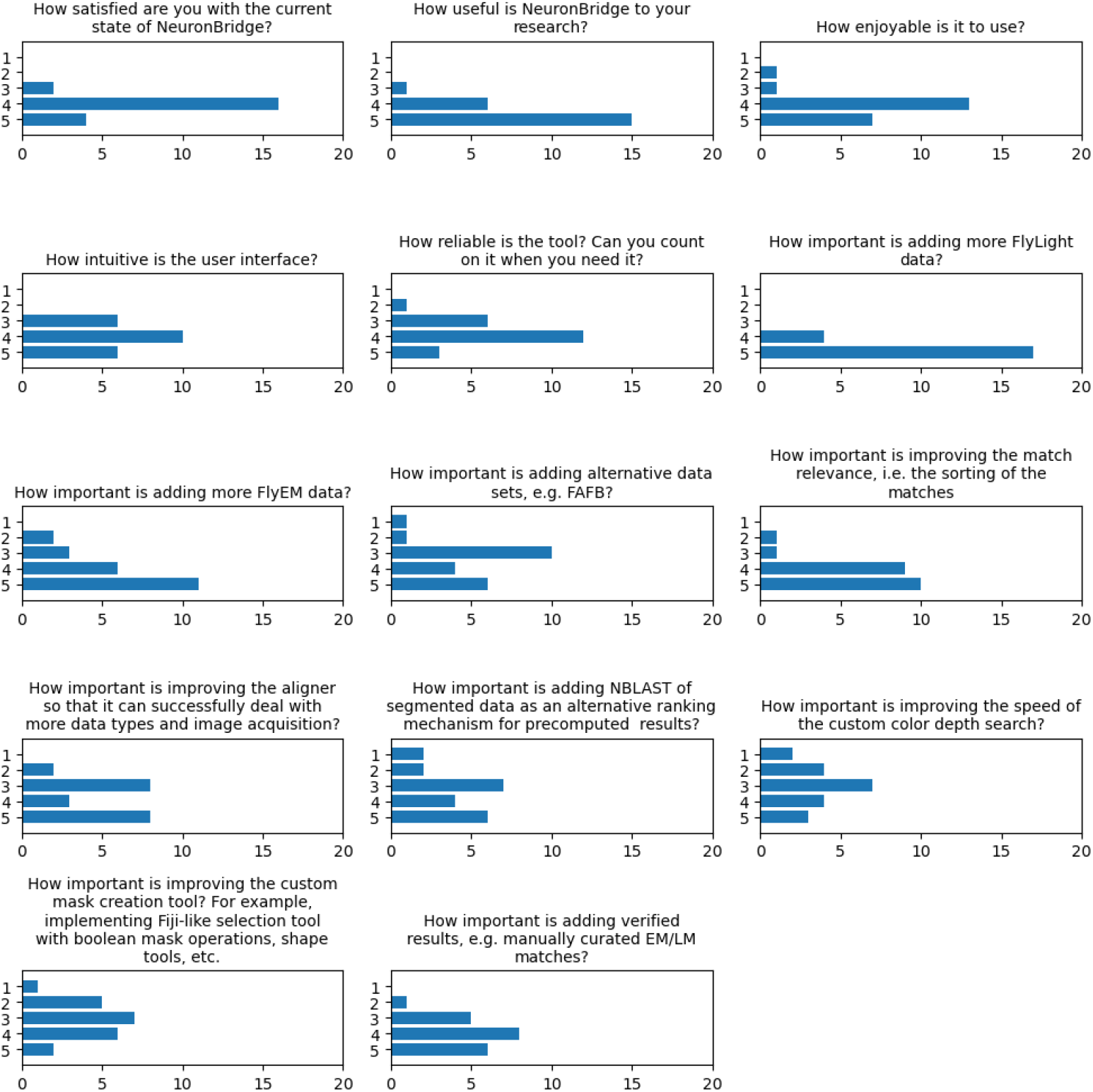
User survey responses. Aggregated responses to the user survey where a value of 1 indicates “Least” and 5 indicates the “Most” response for each question.

**Figure 12.**
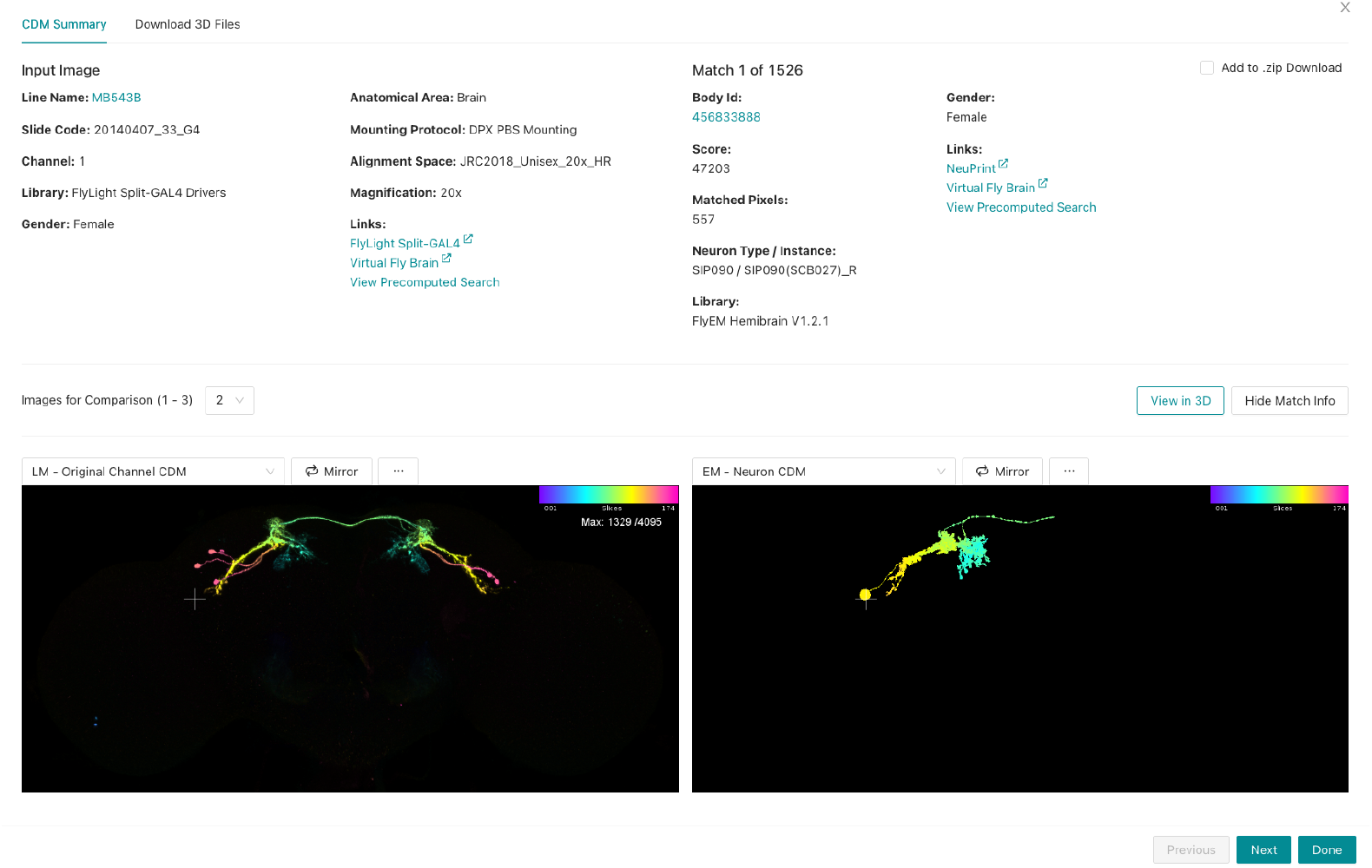
Details for a single EM/LM match. LM input image shown on the left and the EM match on the right. This view is customizable such that the user can choose how many images to display and select which representation to show in each spot.

**Figure 13.**
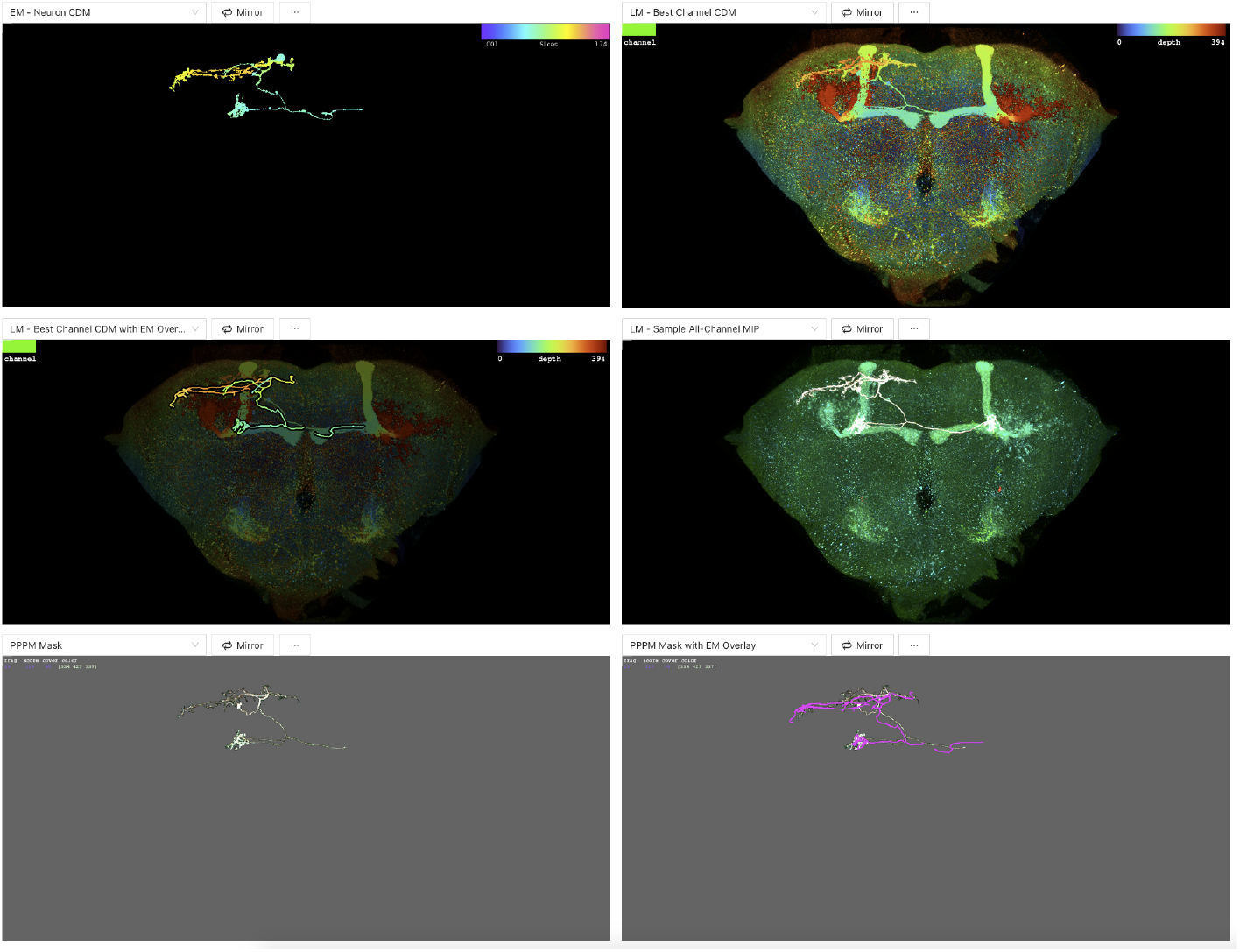
Detail images for a single EM/LM match from PPPM. The PPPM results contain several image types which are not available for CDM matches, including overlays of EM skeletons on masked LM images. These images can be displayed in the configurable image grid.

**Figure 14.**
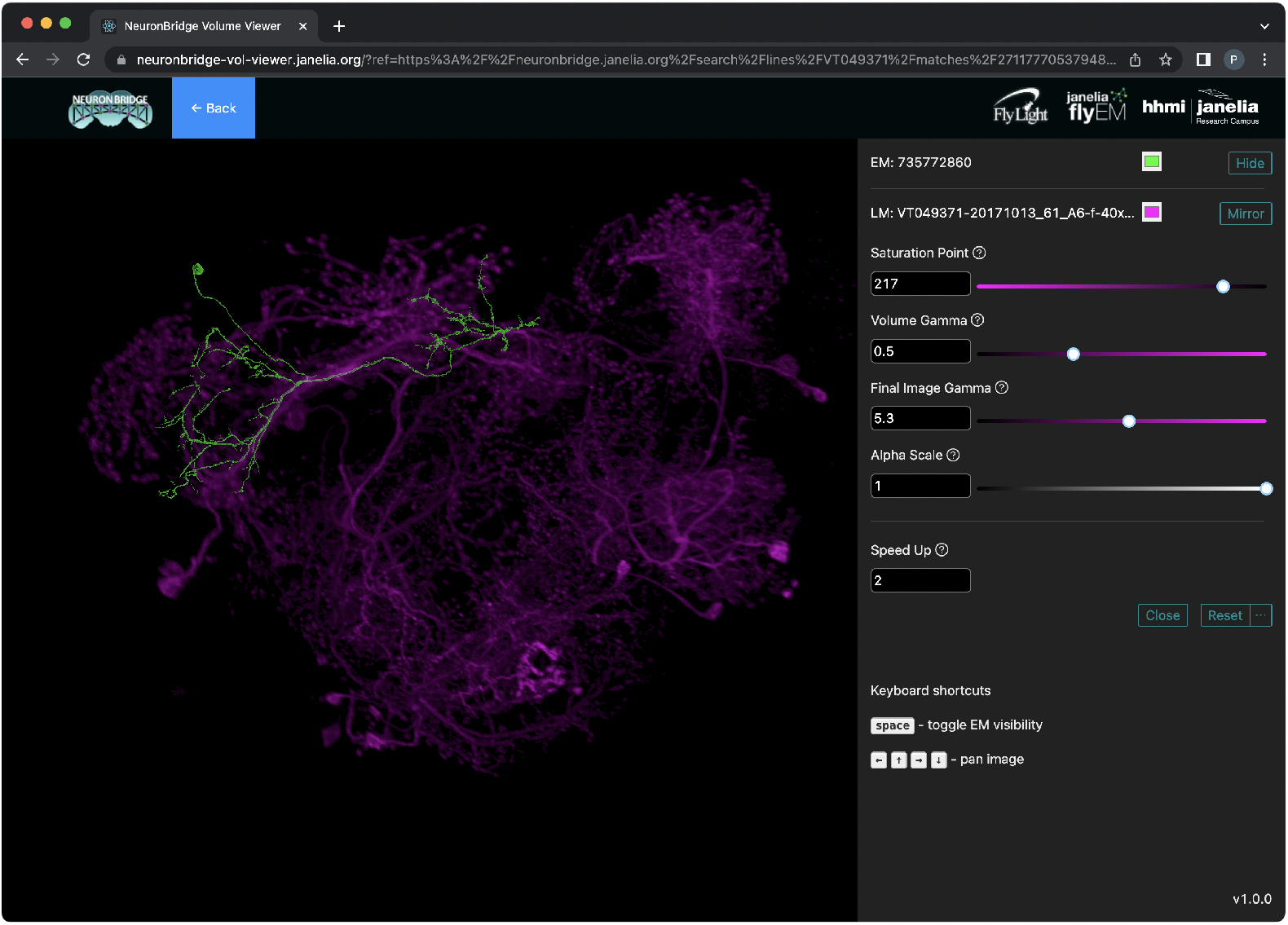
3D visualization of matches in the browser. One channel of the LM volume is rendered in magenta, and the matching EM body is rendered in green. Controls on the right affect qualities of the rendering like the relative transparency in the LM volume.

**Figure 15.**
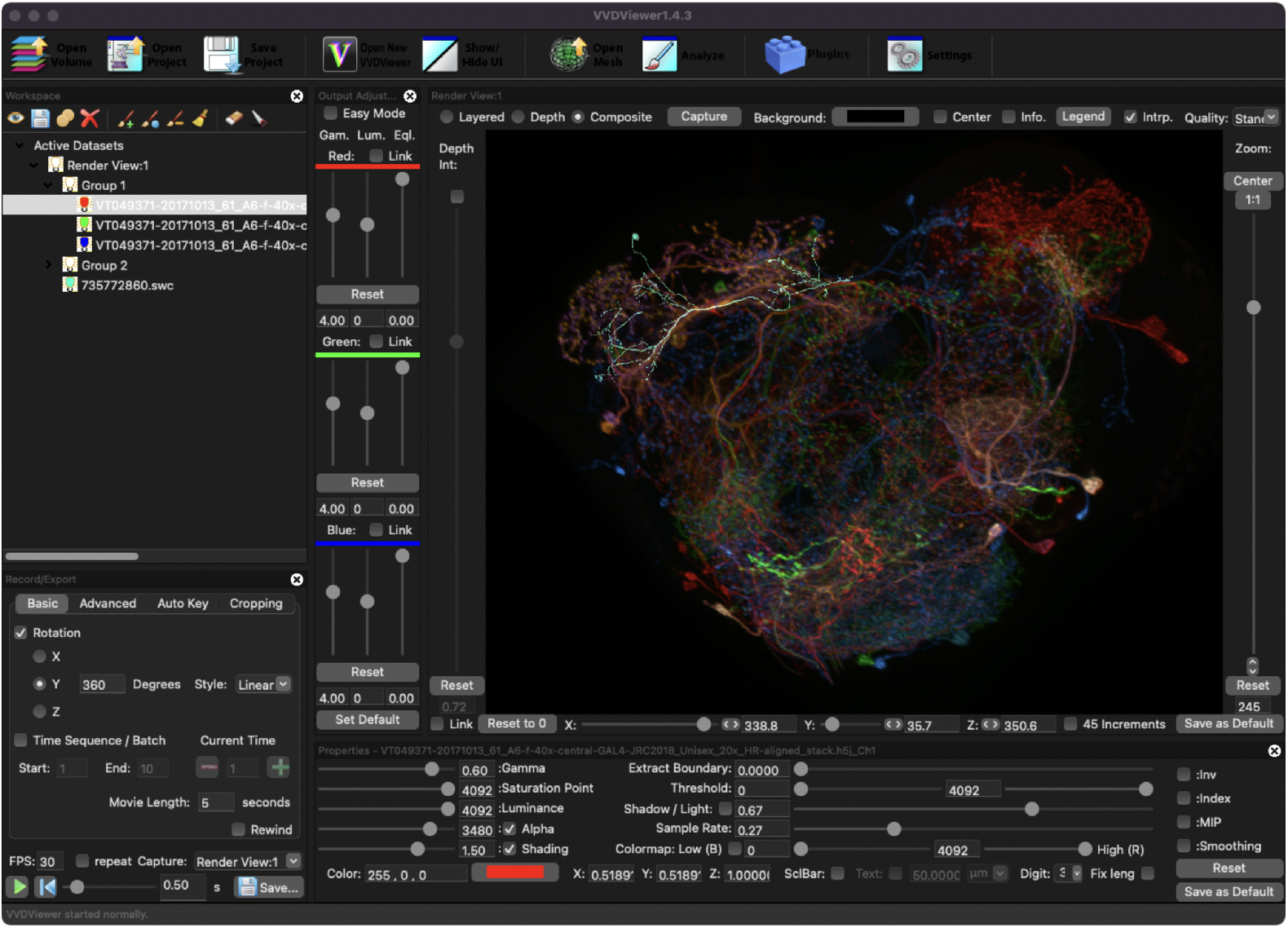
3D visualization of matches in VVD Viewer. All three channels of the LM volume are rendered, in red, green and blue, and the EM body is rendered in aquamarine.

**Figure 16.**
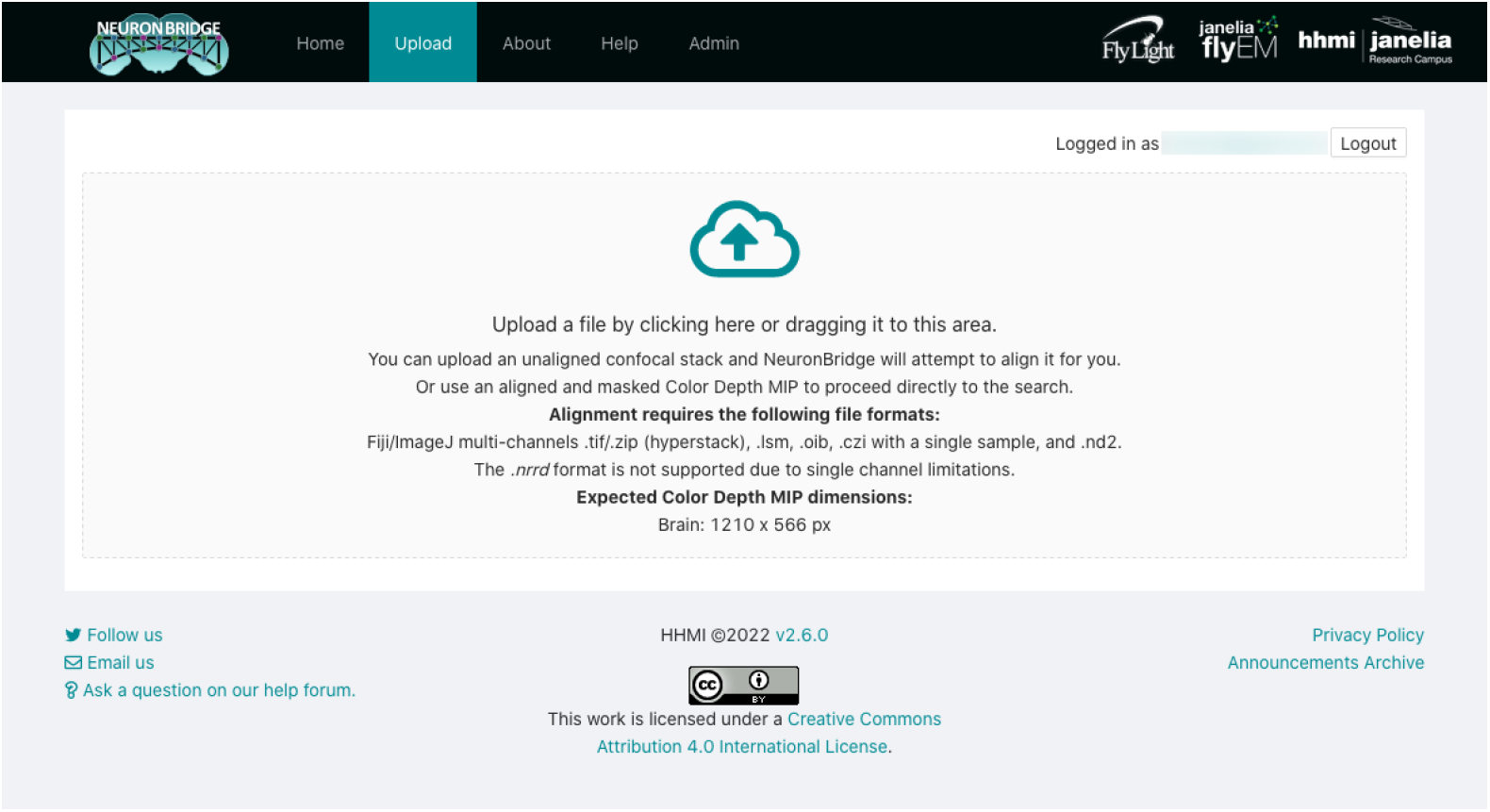
Uploading data for custom search. The upload GUI allows the user to drag and drop a file to begin the custom search workflow. Input data requirements are described in detail.

### Usage Statistics

As of July 2022, there are over one thousand registered users of NeuronBridge. The site received over 28,900 visits in the first half of this year, and 137 user image uploads during that time. During this time period, the API received 8,131 requests from 379 distinct organizations. Since its initial deployment, NeuronBridge has been accessed from 44 countries around the world (Fig. 17).

**Figure 17.**
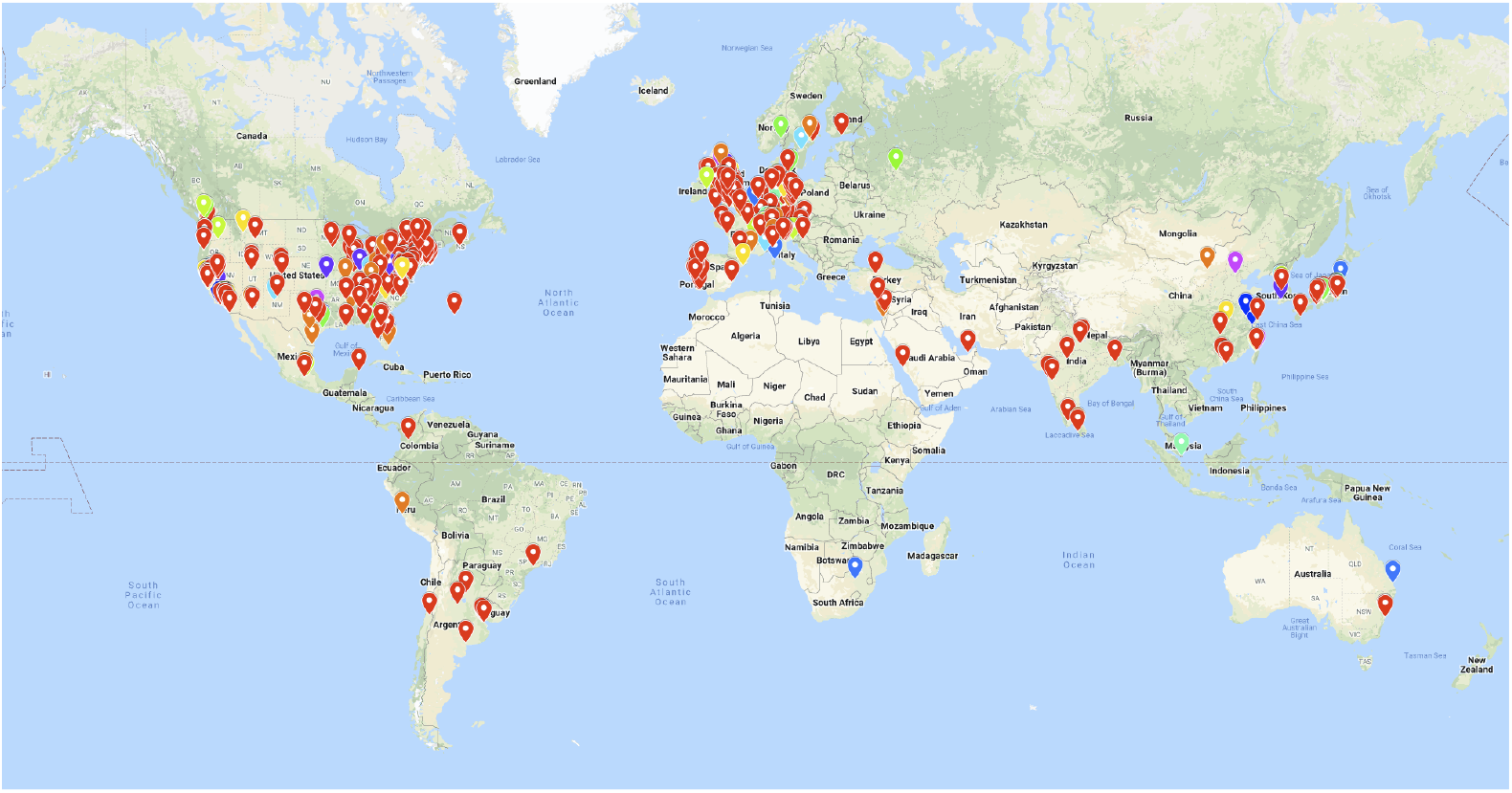
Access map. Depicts access requests to the s3://janelia-neuronbridge-data-prod and s3://janelia-neuronbridge-web-prod buckets during the two year period between 2020-05-08 and 2022-06-08.

## Supplemental Figures

## References

Aso, Yoshinori, Daisuke Hattori, et al. (Dec. 23, 2014). “The neuronal architecture of the mushroom body provides a logic for associative learning”. In: eLife 3. Ed. by Leslie C Griffith, e04577. ISSN: 2050-084X. DOI: 10.7554/eLife.04577.

Aso, Yoshinori and Gerald M Rubin (July 21, 2016). “Dopaminergic neurons write and update memories with cell-type-specific rules”. In: eLife 5. Ed. by Liqun Luo, e16135. ISSN: 2050-084X. DOI: 10.7554/eLife.16135.

Bidaye, Salil S. et al. (Nov. 11, 2020). “Two Brain Pathways Initiate Distinct Forward Walking Programs in Drosophila”. In: Neuron 108.3, 469–485.e8. ISSN: 0896-6273. DOI: 10.1016/j.neuron.2020.07.032.

Bogovic, John, Hideo Otsuna, Shin-ya Takemura, et al. (2022). “Robust search method for Drosophila neurons between electron and light microscopy”. In: (In preparation).

Bogovic, John A., Hideo Otsuna, Larissa Heinrich, et al. (Dec. 31, 2020). “An unbiased template of the Drosophila brain and ventral nerve cord”. In: PLOS ONE 15.12, e0236495. ISSN: 1932-6203. DOI: 10.1371/journal.pone.0236495.

Brand, A. H. and N. Perrimon (June 1993). “Targeted gene expression as a means of altering cell fates and generating dominant phenotypes”. In: Development (Cambridge, England) 118.2, pp. 401–415. ISSN: 0950-1991. DOI: 10.1242/dev.118.2.401.

Clements, Jody et al. (Jan. 17, 2020). neuPrint: Analysis Tools for EM Connectomics, p. 2020.01.16.909465. DOI: 10.1101/2020.01.16.909465.

Costa, Marta et al. (July 20, 2016). “NBLAST: Rapid, Sensitive Comparison of Neuronal Structure and Construction of Neuron Family Databases”. In: Neuron 91.2, pp. 293–311. ISSN: 0896-6273. DOI: 10.1016/j.neuron.2016.06.012.

Davis, Fred P et al. (Jan. 15, 2020). “A genetic, genomic, and computational resource for exploring neural circuit function”. In: eLife 9. Ed. by Hugo J Bellen and K VijayRaghavan, e50901. ISSN: 2050-084X. DOI: 10.7554/eLife.50901.

Dolan, Michael-John et al. (May 21, 2019). “Neurogenetic dissection of the Drosophila lateral horn reveals major outputs, diverse behavioural functions, and interactions with the mushroom body”. In: eLife 8. Ed. by K VijayRaghavan and Ilona C Grunwald Kadow, e43079. ISSN: 2050-084X. DOI: 10.7554/eLife.43079.

Feng, Kai et al. (Dec. 2, 2020). “Distributed control of motor circuits for backward walking in Drosophila”. In: Nature Communications 11.1, p. 6166. ISSN: 2041-1723. DOI: 10.1038/s41467-020-19936-x.

Fouladi, Sadjad et al. (2019). “From Laptop to Lambda: Outsourcing Everyday Jobs to Thousands of Transient Functional Containers”. In: 2019 USENIX Annual Technical Conference (USENIX ATC 19), pp. 475–488. ISBN: 978-1-939133-03-8.

Gao, Shuying et al. (Oct. 23, 2008). “The Neural Substrate of Spectral Preference in Drosophila”. In: Neuron 60.2, pp. 328–342. ISSN: 0896-6273. DOI: 10.1016/j.neuron.2008.08.010.

Hirsch, Peter, Lisa Mais, and Dagmar Kainmueller (Aug. 19, 2020). “PatchPerPix for Instance Segmentation”. In: arXiv:2001.07626 [cs]. arXiv: 2001.07626.

Israel, Shai et al. (Mar. 14, 2022). “Olfactory stimuli and moonwalker SEZ neurons can drive backward locomotion in Drosophila”. In: Current Biology 32.5, 1131–1149.e7. ISSN: 0960-9822. DOI: 10.1016/j.cub.2022.01.035.

Jenett, Arnim et al. (Oct. 25, 2012). “A GAL4-driver line resource for Drosophila neurobiology”. In: Cell Reports 2.4, pp. 991–1001. ISSN: 2211-1247. DOI: 10.1016/j.celrep.2012.09.011.

Kawase, Takashi and Hideo Otsuna (June 28, 2022). “VVD Viewer codebase”. In: GitHub. DOI: 10.5281/zenodo.6762216.

Kawase, Takashi, Shigeo S. Sugano, et al. (Jan. 2015). “A direction-selective local-thresholding method, DSLT, in combination with a dye-based method for automated three-dimensional segmentation of cells and airspaces in developing leaves”. In: The Plant Journal: For Cell and Molecular Biology 81.2, pp. 357–366. ISSN: 1365-313X. DOI: 10.1111/tpj.12738.

Lazar, Aurel A et al. (Feb. 22, 2021). “Accelerating with FlyBrainLab the discovery of the functional logic of the Drosophila brain in the connectomic and synaptomic era”. In: eLife 10. Ed. by Upinder Singh Bhalla, Ronald L Calabrese, and Padraig Gleeson, e62362. ISSN: 2050-084X. DOI: 10.7554/eLife.62362.

Luan, Haojiang et al. (Nov. 9, 2020). “The Drosophila Split Gal4 System for Neural Circuit Mapping”. In: Frontiers in Neural Circuits 14, p. 603397. ISSN: 1662-5110. DOI: 10.3389/fncir.2020.603397.

Mais, Lisa et al. (July 26, 2021). PatchPerPixMatch for Automated 3d Search of Neuronal Morphologies in Light Microscopy, p. 2021.07.23.453511. DOI: 10.1101/2021.07.23.453511.

Meissner, Geoffrey W. et al. (June 1, 2022). A searchable image resource of Drosophila GAL4-driver expression patterns with single neuron resolution. bioRxiv, p. 2020.05.29.080473. DOI: 10.1101/2020.05.29.080473.

Milyaev, Nestor et al. (Feb. 1, 2012). “The Virtual Fly Brain browser and query interface”. In: Bioinformatics 28.3, pp. 411–415. ISSN: 1367-4803. DOI: 10.1093/bioinformatics/btr677.

Morimoto, Mai M et al. (Nov. 18, 2020). “Spatial readout of visual looming in the central brain of Drosophila”. In: eLife 9. Ed. by Claude Desplan, Ronald L Calabrese, and Markus Meister, e57685. ISSN: 2050-084X. DOI: 10.7554/eLife.57685.

Namiki, Shigehiro, Michael H. Dickinson, et al. (Dec. 11, 2017). The functional organization of descending sensory-motor pathways in Drosophila, p. 231696. DOI: 10.1101/231696.

Namiki, Shigehiro, Ivo G. Ros, et al. (Mar. 14, 2022). “A population of descending neurons that regulates the flight motor of Drosophila”. In: Current Biology 32.5, 1189–1196.e6. ISSN: 0960-9822. DOI: 10.1016/j.cub.2022.01.008.

Nern, Aljoscha, Barret D. Pfeiffer, and Gerald M. Rubin (June 2, 2015). “Optimized tools for multicolor stochastic labeling reveal diverse stereotyped cell arrangements in the fly visual system”. In: Proceedings of the National Academy of Sciences 112.22, E2967–E2976. ISSN: 0027-8424, 1091-6490. DOI: 10.1073/pnas.1506763112.

Nojima, Tetsuya et al. (Mar. 22, 2021). “A sex-specific switch between visual and olfactory inputs underlies adaptive sex differences in behavior”. In: Current Biology 31.6, 1175–1191.e6. ISSN: 0960-9822. DOI: 10.1016/j.cub.2020.12.047.

Otsuna, Hideo, Masayoshi Ito, and Takashi Kawase (May 9, 2018). Color depth MIP mask search: a new tool to expedite Split-GAL4 creation, p. 318006. DOI: 10.1101/318006.

Pfeiffer, Barret et al. (Oct. 1, 2010). “Refinement of Tools for Targeted Gene Expression in Drosophila”. In: Genetics 186, pp. 735–55. DOI: 10.1534/genetics.110.119917.

Robie, Alice A. et al. (July 13, 2017). “Mapping the Neural Substrates of Behavior”. In: Cell 170.2, 393–406.e28. ISSN: 0092-8674, 1097-4172. DOI: 10.1016/j.cell.2017.06.032.

Rohlfing, Torsten and Calvin R. Maurer (Mar. 2003). “Nonrigid image registration in shared-memory multiprocessor environments with application to brains, breasts, and bees”. In: IEEE transactions on information technology in biomedicine: a publication of the IEEE Engineering in Medicine and Biology Society 7.1, pp. 16–25. ISSN: 1089-7771. DOI: 10.1109/titb.2003.808506.

Rokicki, Konrad (Feb. 10, 2021). Scaling Neuroscience Research on AWS. AWS Architecture Blog. URL: https://aws.amazon.com/blogs/architecture/scaling-neuroscience-research-on-aws/.

Sareen, Preeti F., Li Yan McCurdy, and Michael N. Nitabach (July 5, 2021). “A neuronal ensemble encoding adaptive choice during sensory conflict in Drosophila”. In: Nature Communications 12.1, p. 4131. ISSN: 2041-1723. DOI: 10.1038/s41467-021-24423-y.

Scheffer, Louis K et al. (Sept. 3, 2020). “A connectome and analysis of the adult Drosophila central brain”. In: eLife 9. Ed. by Eve Marder et al., e57443. ISSN: 2050-084X. DOI: 10.7554/eLife.57443.

Schindelin, Johannes et al. (July 2012). “Fiji: an open-source platform for biological-image analysis”. In: Nature Methods 9.7, pp. 676–682. ISSN: 1548-7105. DOI: 10.1038/nmeth.2019.

Schlegel, Philipp et al. (May 25, 2021). “Information flow, cell types and stereotypy in a full olfactory connectome”. In: eLife 10. Ed. by Leslie C Griffith, Catherine Dulac, and Liqun Luo, e66018. ISSN: 2050-084X. DOI: 10.7554/eLife.66018.

Schretter, Catherine E et al. (Nov. 3, 2020). “Cell types and neuronal circuitry underlying female aggression in Drosophila”. In: eLife 9. Ed. by Mani Ramaswami and Catherine Dulac, e58942. ISSN: 2050-084X. DOI: 10.7554/eLife.58942.

Sterne, Gabriella R et al. (Sept. 2, 2021). “Classification and genetic targeting of cell types in the primary taste and premotor center of the adult Drosophila brain”. In: eLife 10. Ed. by Ilona C Grunwald Kadow, Ronald L Calabrese, and Kristen Lee, e71679. ISSN: 2050-084X. DOI: 10.7554/eLife.71679.

Tanaka, Ryosuke and Damon A. Clark (May 3, 2022). “Neural mechanisms to exploit positional geometry for collision avoidance”. In: Current Biology. ISSN: 0960-9822. DOI: 10.1016/j.cub.2022.04.023.

Tuthill, John C. et al. (July 10, 2013). “Contributions of the 12 Neuron Classes in the Fly Lamina to Motion Vision”. In: Neuron 79.1, pp. 128–140. ISSN: 0896-6273. DOI: 10.1016/j.neuron.2013.05.024.

Wang, Kaiyu et al. (Jan. 2021). “Neural circuit mechanisms of sexual receptivity in Drosophila females”. In: Nature 589.7843, pp. 577–581. ISSN: 1476-4687. DOI: 10.1038/s41586-020-2972-7.

Wolff, Tanya and Gerald M. Rubin (2018). “Neuroarchitecture of the Drosophila central complex: A catalog of nodulus and asymmetrical body neurons and a revision of the protocerebral bridge catalog”. In: Journal of Comparative Neurology 526.16, pp. 2585–2611. ISSN: 1096-9861. DOI: 10.1002/cne.24512.

Wu, Ming et al. (Dec. 28, 2016). “Visual projection neurons in the Drosophila lobula link feature detection to distinct behavioral programs”. In: eLife 5. Ed. by Kristin Scott, e21022. ISSN: 2050-084X. DOI: 10.7554/eLife.21022.

Xu, C Shan et al. (May 13, 2017). “Enhanced FIB-SEM systems for large-volume 3D imaging”. In: eLife 6. Ed. by Jeremy Nathans, e25916. ISSN: 2050-084X. DOI: 10.7554/eLife.25916.

